# Unbalanced Sample Size Introduces Spurious Correlations to Genome-wide Heterozygosity Analyses

**DOI:** 10.1101/2020.02.06.937599

**Authors:** Li Liu, Richard J Caselli

## Abstract

Excess of heterozygosity (H) is a widely used measure of genetic diversity of a population. As high-throughput sequencing and genotyping data become readily available, it has been applied to investigating the associations of genome-wide genetic diversity with human diseases and traits. However, these studies often report contradictory results. In this paper, we present a meta-analysis of five whole-exome studies to examine the association of H scores with Alzheimer’s disease. We show that the mean H score of a group is not associated with the disease status, but is associated with the sample size. Across all five studies, the group with more samples has a significantly lower H score than the group with fewer samples. To remove potential confounders in empirical data sets, we perform computer simulations to create artificial genomes controlled for the number of polymorphic loci, the sample size and the allele frequency. Analyses of these simulated data confirm the negative correlation between the sample size and the H score. Furthermore, we find that genomes with a large number of rare variants also have inflated H scores. These biases altogether can lead to spurious associations between genetic diversity and the phenotype of interest. Based on these findings, we advocate that studies shall balance the sample sizes when using genome-wide H scores to assess genetic diversities of different populations, which helps improve the reproducibility of future research.

## Introduction

Genetic diversity concerns the amount of standing genetic variations in an individual genome or in a population [1]. Across species and evolutionary time spans, populations with a high genetic diversity often have higher fitness than those with a low genetic diversity, likely due to enhanced adaptability to environmental changes [2-4]. For example, heterozygous advantage has been observed in lab settings and in clinical settings. Mammals that are heterozygous at the major histocompatibility complex locus are more resistant to multiple-strain infections than homozygotes [5]. Cells that are heterozygous at the triosephosphate isomerase locus respond to oxidative stresses better than homozygotes [6]. These findings have significant implications for health management, species conservation and other biomedical fields.

Recently, as high-throughput sequencing and genotyping data become increasingly accessible, studies have emerged that expand diversity analyses to the genomic scale and examine the associations with human phenotypes (reviewed in [7-9]). However, these studies often produce contradictory results [10]. For instance, Campbell et al. reported significant associations of low genome-wide heterozygosity with high blood pressures [11], while three other studies did not observe these relationships [12-14]. Similarly, Bihlmeyer et al. and Xu et al. independently reported positive impact of genomic diversity on human mortality and longevity [15, 16]. However many studies of non-human species have shown that such impact is only significant in extremely harsh environments [17-19], which raises the question of the mechanisms in human populations. Furthermore, some reported associations between genetic diversity and human phenotypes vanished when including additional samples in the analyses [20, 21].

Many factors may contribute to the lack of reproducibility of these studies, such as population stratifications, sequencing and genotyping platforms, clinical and demographic cofounders, and statistical approaches. Here, we examine a specific measure, i.e., excess of heterozygosity (H) that is commonly employed to quantify and compare genetic diversity in different populations. The H score was originally developed in the pre-genomic era to estimate genetic diversity at a small number of polymorphic loci and to detect population bottlenecks in a single set of samples [22, 23]. Although its extension to include tens of thousands of variants from multiple phenotypically different groups is computationally intuitive, the influences of these adaptations on statistical inferences remain unclear. Current studies have used the H score blindly and paid little attention to potential pitfalls. To address these questions, we perform a meta-analysis of associations between exome-wide H scores with Alzheimer’s disease (AD). We then investigate the impact of the number of polymorphic loci, the allele frequency and the number of samples on H score estimations and subsequent association statistics. We further conduct computer simulations to confirm the findings and propose a sample balancing strategy to rectify the potential biases.

## Materials and Methods

### Processing of AD data sets

The Alzheimer’s Disease Sequencing Project (ADSP) [24] is a multi-institutional effort to collect and analyze whole-genome and whole-exome data for AD research. Via the dbGaP portal (accession number phs000572.v7.p4.c1-6), we queried the meta-data of the ADSP to identify cases diagnosed with late-onset AD (estimated onset age ≥65) and elder controls (cognitively normal at age ≥80) from five independent cohorts, namely the ACT, ADC, CHS, LOAD and RS cohorts. We then downloaded the whole-exome data of these samples in the VCF format and used the vcftools [25] to extract bi-allelic single-nucleotide variants (SNVs) in autosomes with minor allele frequencies (MAF) >0.1%. We set the minimum genotyping quality to 20 and removed sites violating the Hardy-Weinberg equilibrium (p-value <0.05).

### Compute excess of heterozygosity

For each study, we computed an H score at each bi-allelic site among cases and among controls, separately. Given a group of samples genotyped at a specific site harboring a major allele and a minor allele, the observed heterozygosity rate *O* is the fractions of heterozygous samples over the total samples. The expected heterozygosity rate is *E* = 1 – *f*^2^ – (1 – *f*)^2^ where f is the MAF. The excess of heterozygosity is 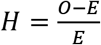. Large H scores indicate high genetic diversity.

### Statistical Analysis

To compare H scores between two groups, we used two-sided Student’s t test. To test associations between H scores and other controlling factors, we used linear regression models. We applied the Bonferroni correction to adjust the p-values for multiple comparisons.

### Computer simulations

To remove potential confounders, we created artificial genomes in which all variants were random samples from the same population with a given MAF. At a given site, we assumed that the major and minor alleles followed a binomial distribution *B(n, p)* where *n* was set to the population size and *p* was set to the MAF value, and simulated genotypes accordingly. From these artificial genomes, we drew two random subsets to represent two groups of samples. We then computed H scores for each polymorphic site within each group of samples and used t-test to assess if the mean H scores were significantly different between the two groups. We varied the MAF value between 0.1% and 0.5, the number of polymorphic loci between 10 and 5,000, and the sample size between 20 and 1,000. For each combination of these parameter values, we generated 10 simulations. In total, we produced 11,340 simulations.

## Results

### Meta-analysis of AD studies

We obtained whole-exome sequences for 2,942 late-onset AD cases and 1,705 elder non-AD controls from five independent cohorts (**Table 1**). All cohorts involved unequal numbers of AD samples and non-AD samples. There were more non-AD samples than AD samples in the ACT, CHS and RS cohorts, and more AD samples than non-AD samples in the ADC and LOAD cohorts. Because APOE genotype is a known risk factor for AD [26], we also stratified samples into an e3/e3 subset and an e3/e4 subset. On average, each AD study reported 48,681 SNVs with MAF>0.01 in autosomes (range: 43,675 to 55,611).

**Table 1.** Summary of samples in each AD study.

| Study Name | All Samples (N) | AD Cases * (N<sub>AD</sub>) | Non-AD Controls * (N<sub>non-AD</sub>) | Total SNVs |
| --- | --- | --- | --- | --- |
| ACT | 1,181 | 285 (230 \| 17) | 896 (610 \| 131) | 45,234 |
| ADC | 1,421 | 768 (525 \| 157) | 653 (430 \| 85) | 55,611 |
| CHS | 788 | 225 (180 \| 14) | 563 (391 \| 51) | 43,675 |
| LOAD | 256 | 181 (73 \| 95) | 75 (52 \| 0) | 55,157 |
| RS | 1,001 | 246 (182 \| 33) | 755 (472 \| 116) | 43,728 |

\* Numbers of samples with the e3/e3 and the e3/e4 APOE genotypes are shown inside parentheses.

Within a study, we computed an H score at each SNV locus for the AD group and for the non-AD group separately. We then performed t-tests to compare the mean H scores between the two groups. In three cohorts, namely ACT, CHS and RS, the AD group had a significantly higher mean H score than the non-AD group (all p-value < 0.01, **Table 2**). However, the other two studies showed the reverse trend. Interestingly, the conversion of the relative H scores coincided with the relative sample sizes of the AD and non-AD groups (**Fig. 1A**). In studies where AD samples outnumbered non-AD samples, the mean H scores of the AD group was lower than that of the non-AD group (Pearson correlation coefficient = −0.90, *P* = 0.036). This negative correlation persisted when we stratified the samples based on their APOE genotypes and performed the analyses within each stratum (**Fig. 1B-C**).

**Table 2.**
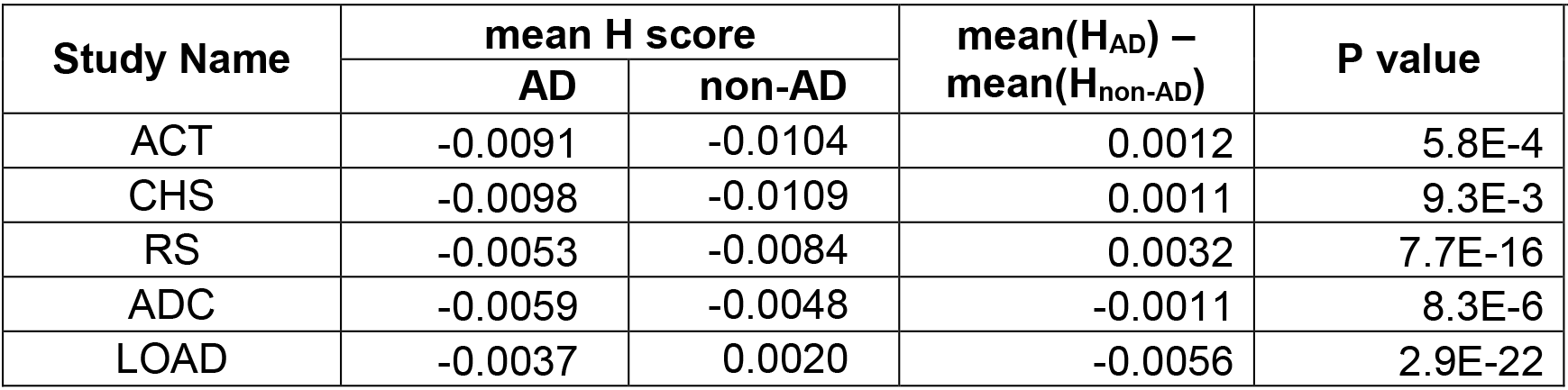
Comparison of mean H scores between AD and non-AD groups

**Figure 1.**
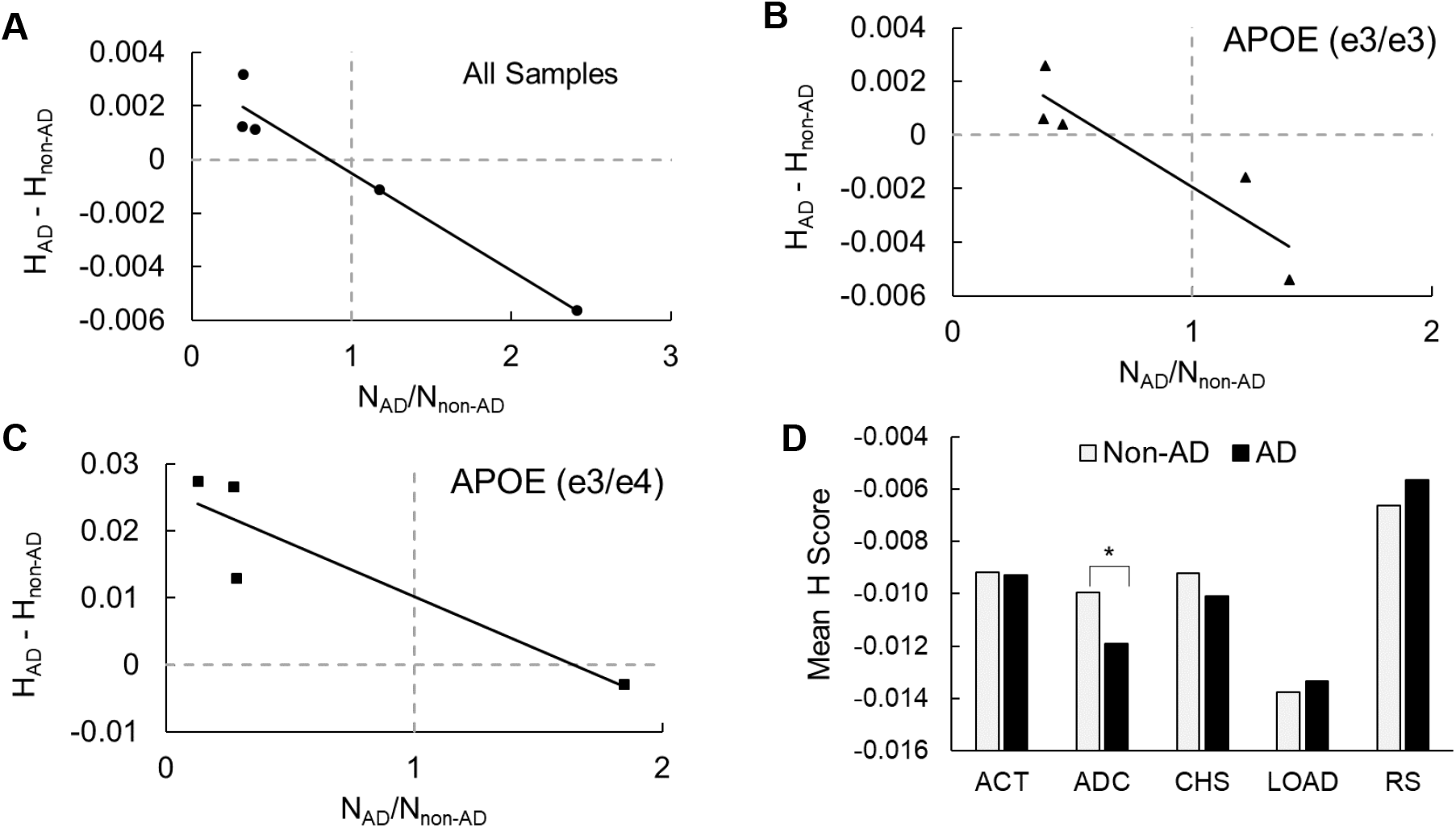
Meta-analysis of five AD cohorts. (**A, B, C**) Scatterplots show the relationship of sample size imbalance with mean difference of H scores between the AD group and the non-AD group. For each cohort, all samples (A), samples with the APOE e3/e3 genotype (B), and samples with the APOE e3/d4 genotype (C) are used in the analyses. Solid black lines represent linear fits. Vertical dotted lines represent equal sample sizes. Horizontal dotted lines represent equal H scores. (**D**) A bar plot shows mean H scores of the AD group and of the non-AD group in each cohort after balancing the sample sizes. The difference is only significant in the ADC cohort, as marked by the asterisk.

We then down-sampled the large group to the same size of the small group in each study, and repeated the analysis. In four of the five cohorts, we observed no significant differences in H scores between the AD group and the non-AD group (all *P* values > 0.05, **Fig. 1D**). The only exception was the ADC cohort where the mean H score of the non-AD group was higher than that of the AD group (−0.010 vs. −0.012, *P*=2.4×10^−7^).

These results implied that unbalanced sample size instead of disease status likely contributed to the differences in H scores between the AD group and the non-AD group. However, other factors, such as sex, disease subtypes and sequencing procedures might have confounded the analyses. Because it is not feasible to measure and include all these factors in the analysis of empirical data, we next performed computer simulations with strictly controlled parameters.

### Computer simulations and analyses

In each simulation, we created a population of 1 million artificial genomes. All artificial genomes in this population contained the same number of polymorphic loci, and the major and minor alleles at each locus followed the same binomial distribution defined by the population size and a specific MAF. We then randomly drew two sets of genomes from this population, which should have no differences in genetic diversity. If the Bonferroni-corrected t-test P value indicated significantly different H scores between the two sets, we considered it as a false positive association. We varied the number of polymorphic loci (*L*), the MAF (*F*) and the sample size (*N1 and N2* with *N1 ≤ N2*) of each random set to simulate different scenarios. For each unique combination of these four parameters, we produced 10 simulations, and recorded the fraction of simulations giving false positive results (i.e., the false positive rate, *FPR*).

Among the total of 11,340 simulation, 3,323 reported false positive associations. In all of these false positive cases, the small group had a higher H score than the large group, confirming the negative correlation between the sample size and the H score (Fisher’s exact test *P*<10^−16^).

We then used a linear regression model to examine the relationship between *FPRs* and the four controlled parameters (**Table 3**). This model showed that FPRs increased when H scores were computed from more polymorphic loci (*P*=10^−16^) harboring low-frequency variants (*P*=0.004), which mimicked the scenarios in high-throughput sequencing studies. *FPR*s also increased as the imbalance of sample sizes, represented by the *N2/N1* ratio, became more severe (*P*=10^−14^). Due to the combined effects of the three factors, when less than 1,000 loci were present, the H scores were robust to sample size imbalance (**Fig. 2A-B**). In contrast, when more than 10,000 loci were present, even a small sample size imbalance could lead to high FPRs (**Fig. 2C-D**).

**Table 3.** Linear regression model of FPR against control parameters

| Control Parameter | Coefficient | P value |
| --- | --- | --- |
| F | -1.4E-01 | 4.2E-03 ** |
| N1 | -4.1E-04 | 6.2E-15 *** |
| N2 | 6.4E-05 | 6.5E-02 |
| N2/N1 | 8.3E-03 | 1.1E-14 *** |
| L | 6.6E-06 | 2.0E-16 *** |

**Figure 2.**
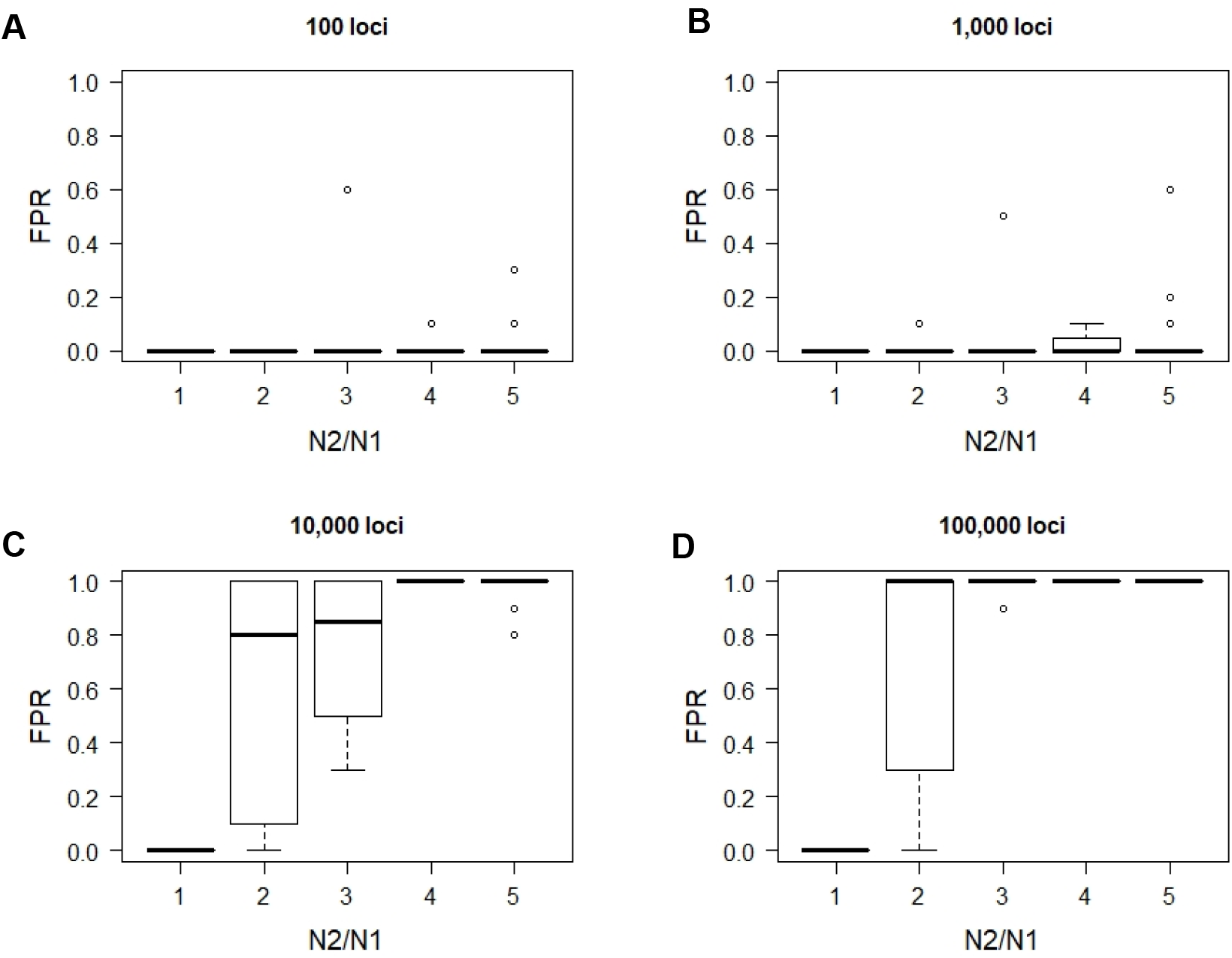
False positive rates (FPRs) increase with sample size imbalance. Boxplots show the distributions of FPRs when different numbers of loci are used to compute H scores. For each N2/N1 ratio, variants of different MAFs are aggregated.

Lastly, consistent with the general statistical principles, we found that larger sample sizes reduced FPRs (*P*=10^−15^).

## Discussion/Conclusion

Studies have emerged that investigate the role of genome-wide genetic diversity in human diseases and traits [11-16]. These studies usually adapt traditional population genetic measures, such as excess of heterozygosity [27], runs of homozygosity [28] and inbreeding coefficients [29] that were originally developed to measure diversity at a small number of genetic loci to analyzing tens of thousands of loci produced from high-throughput sequencing data. In this study, we focused on the excess of heterozygosity score and examined the potential biases introduced by the escalation of scales of analysis.

Our most important discovery is the dependency of H scores on sample sizes. When comparing H scores between two groups of samples, the larger group tends to produce lower H scores than the smaller group. Such biases are more severe when H scores are computed from a large number of rare variants. Unfortunately, these are common characteristics of data from whole-genome and whole-exome disease association studies, in which tens of thousands to millions of genetic variants are identified and little attention is paid to balance the number of cases and controls. Such biases contribute at least partially to the inconsistent results from existing studies.

In our meta-analysis of whole-exome data from five AD cohorts, we indeed observed that the difference of H scores between the AD and non-AD groups coincided with the relative sample size of these two groups. In the three cohorts where non-AD samples outnumbered AD samples, high genetic diversity was a significant risk factor. In contrast, in the two cohorts where AD samples outnumbered non-AD samples, high genetic diversity became a protective factor. However, these associations vanished after we balanced the sample sizes. The only exception was the ADC cohort, in which high genetic diversity was a protective factor both before and after sample size balancing. Given these results and the high heterogeneity of AD, it is premature to infer an association (or lack thereof) of exome-wide H scores and AD risk at the current stage.

A straightforward solution to correcting these biases is to ensure each group contains the same number of samples. In addition to simple balanced sub-sampling, genomic matching is another approach, in which the genomic profile of a sample from one phenotypic group is used to find the best-matched sample in the other groups [15]. Genomic matching achieves balanced sample sizes and minimal between-group genomic distances simultaneously. When the association involves quantitative traits, stratified balancing can be applied that first discretizes the quantitative trait and then matches samples within each stratum [30].

Our findings help to explain the conflicting results from existing studies and will hopefully improve the reproducibility of future research.

## Acknowledgement

We thank Dr. David Brafman at Arizona State University and Drs. Matt Huentelman and Ignazio Piras at the Translational Genomics Research Institute for insightful discussions. We thank the ADSP and dbGaP for providing genomic and clinical data used in this study.

## Author Contributions

LL and RJC designed the study. LL conducted the analysis. Both authors wrote the manuscript.

